# BiomniBench: Process-level Evaluation of LLM Agents for Real-world Biomedical Research

**DOI:** 10.64898/2026.05.12.724604

**Authors:** Yuanhao Qu, Yingzhou Lu, Xinming Tu, Serena Zhang, Tianwei She, Alexander Glenn Shaw, Jou-Ho Shih, Bingqing Zhao, Minjie Shen, Haochen Yang, Jielin Yan, Rongchuan Zhang, Xinze Wu, Tingting Li, Bin Zhou, Ning Wang, Adam Ma, Le Cong, Xiaobo Hu, Yuan Jiang, Jiayun Dong, Tao Peng, Jure Leskovec, Kexin Huang

## Abstract

LLM agents now perform real biomedical research, but evaluating them rigorously is hard. Outcome-only benchmarks fail in two ways. First, a correct final answer can come from memorization, reward hacking, or wrong reasoning that produces the right number by chance. Second, valid alternative analyses are marked wrong simply because they differ from the reference. We introduce BiomniBench, a process-level evaluation framework that scores the full agent trajectory against expert-designed, task-specific rubrics. Our first release, BiomniBench-DA, contains 100 data-analysis tasks across 17 task types, 5 disease areas, and a general-biology category, each based on a paper from journals such as Nature, Cell, and Science and co-developed with an original author or a domain expert. Benchmarking frontier and open-weight models across four agent harnesses reveals three findings. Frontier and open-weight bases cluster within a few points of each other, with substantial headroom for all models. The agent harness shifts scores by more than the gap between successive model generations. Agents reliably ground claims in real sources yet consistently fall short on method selection, biological interpretation, and scientific reasoning. BiomniBench is the first process-level benchmark for LLM agents in biomedical research, providing the dimension-level diagnostics that outcome scoring cannot.

**Dataset:** huggingface.co/datasets/phylobio/BiomniBench-DA

## 1 Introduction

Large language model (LLM) agents are reshaping biological research. Combining language understanding with code execution, tool use, and structured database access, today’s agents can design CRISPR and genetic-perturbation experiments, analyze spatial transcriptomic data, plan autonomous chemistry workflows, support drug-discovery pipelines, and interpret clinical multi-omics — work that once required hours to days of expert effort [Huang et al., 2025, Qu et al., 2026, Roohani et al., 2025, Boiko et al., 2023, M. Bran et al., 2024, Liu et al., 2024, Wang et al., 2025]. As these capabilities have matured, both specialized biomedical agents and general-purpose coding agents such as Claude Code and OpenAI Codex are being seriously adopted into real scientific workflows across academia and industry, raising a question: how do we know when these agents are actually doing good science?

Most existing benchmarks try to answer this with final-answer scoring: exact match, binary correctness, or pass/fail on a held-out result, e.g., HLE [Phan et al., 2025] and BixBench [Mitchener et al., 2025]. This works for problems with a single verifiable answer, but for complex biomedical research, outcome-only evaluation fails in two ways. First, final-answer benchmarks invite data contamination and reward hacking: published datasets often appear in training corpora, and outcome scoring rewards landing on the answer rather than doing the analysis. An agent can return a clean volcano plot, a ranked gene list, and a confident interpretation while its trace reveals wrong normalization, ignored batch effects, and a fabricated citation. Second, real research is open-ended: multiple analytical paths yield different but defensible answers, so outcome-matching penalizes alternatives that diverge from the reference even when the analysis is sound. In biology these failures are especially costly, because flawed analyses propagate silently into wet-lab experiments, drug-development decisions, and downstream research pipelines for months before anyone notices.

We introduce **BiomniBench**, a process-level evaluation framework for LLM agents on biomedical research tasks. BiomniBench grades the full agent trajectory against expert-designed, task-specific rubrics applied by an LLM judge, rather than scoring only the final result (Figure 1). Three design principles shape the framework. **(1) Ground tasks in real-world research**. Every task is derived from published research and requires multi-step reasoning, method selection, and interpretation. **(2) Evaluate the process, not just the output**. Rubrics score the quality of analytical decisions at each step, distinguishing flawed methods that land on the right answer from rigorous methods that miss by a small margin. **(3) Accommodate multiple valid approaches**. Each rubric encodes alternative analytical paths so agents are not penalized for methodological choices that differ from the reference analysis.

**Figure 1:**
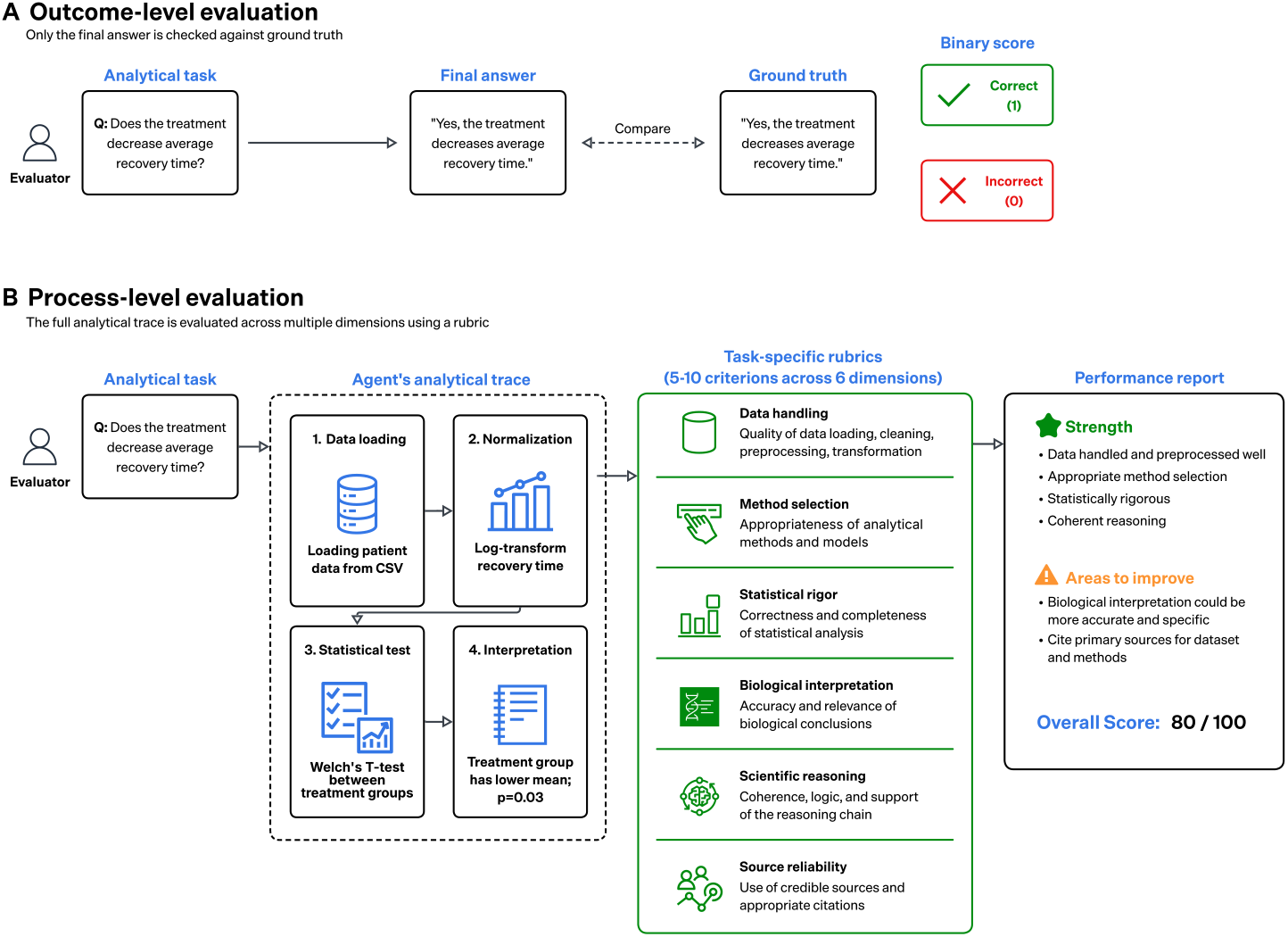
Process-level vs. outcome-level evaluation. (A) Outcome-only evaluation compares only the agent’s final output to a reference answer. (B) Process-level evaluation scores the full agent trajectory against a task-specific rubric, producing dimension-level diagnostics that pinpoint where the agent fails.

To choose the framework’s first release, we analyzed 32,014 user-submitted tasks from the opensource Biomni platform [Huang et al., 2025]; details in Appendix D. Data analysis dominates at 63.3%, well ahead of literature research, experimental design, quick factual queries, and manuscript writing. We therefore release **BiomniBench-DA**: 100 data analysis tasks curated from papers in Nature, Cell, Science, and similar journals, spanning 17 task types across 5 disease areas and a general-biology category. Each task is co-developed with either an original paper author or a domain expert with 5+ years of research experience, and includes the underlying public dataset, a multi-step reference trace, and a task-specific rubric. The rubric scores six dimensions: data handling, method selection, statistical rigor, biological interpretation, scientific reasoning, and source reliability.

We benchmark LLMs across four agent harnesses (Codex CLI, Terminus-2, Claude Code, Gemini CLI), spanning closed-source frontier systems (Claude Opus 4.7, GPT-5.5, Gemini 3.1 Pro) and open-weight models (GLM-5.1, Qwen 3.6, Kimi K2.6). Frontier and open-weight bases cluster within a few points of each other, with substantial headroom for all models. The agent harness shifts scores by more than the gap between successive model generations. Dimension-level analysis reveals that agents reliably ground claims in real sources yet consistently fall short on method selection, biological interpretation, and scientific reasoning. Together, these results establish BiomniBench as the first process-level benchmark for LLM agents on real-world biomedical research, giving model developers, agent builders, and the scientific community a shared benchmark for tracking progress on capabilities that outcome-only evaluation cannot measure.

## 2 BiomniBench-DA: Benchmark design

BiomniBench-DA presents agents with biomedical research tasks as researchers encounter them: a published study supplies the question and reference solution, a rubric co-developed with the paper authors grades the agent’s trace, and a standardized environment inside the Harbor execution framework [Harbor Framework Team, 2026] unifies outputs across harnesses.

### 2.1 Expert-driven curation pipeline

The 100 BiomniBench-DA tasks are produced by a five-stage pipeline that begins with an analysis of biomedical-agent usage and ends with a task-specific rubric (Figure 2, top).

**Figure 2:**
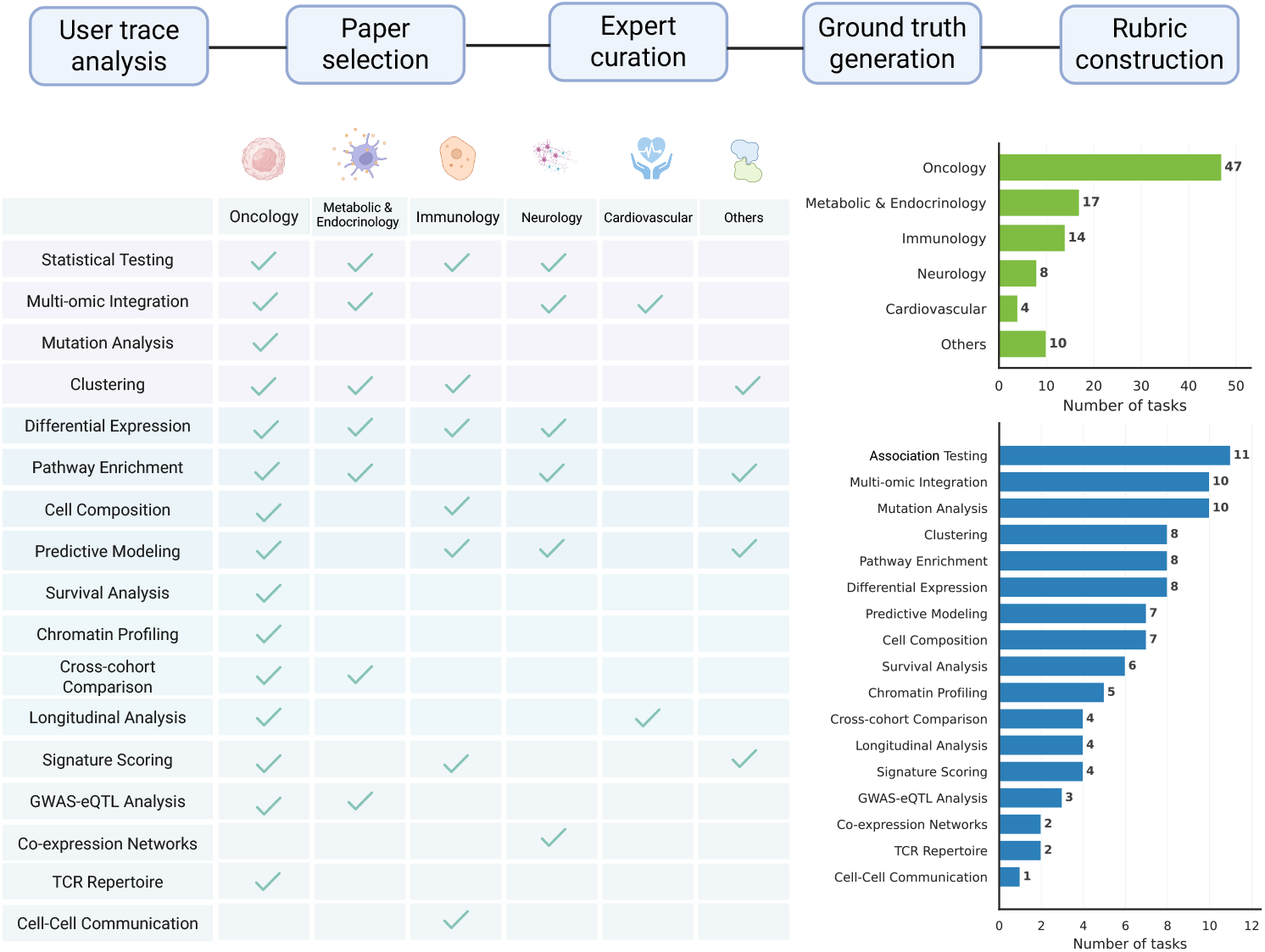
BiomniBench-DA overview (n = 100 tasks). **(Top)** Five-stage curation pipeline: user-trace analysis, paper selection, expert curation, ground-truth generation, and rubric design. **(Bottomleft)** Coverage matrix of (task type, disease category) combinations; a checkmark indicates that the benchmark includes at least one task at that intersection. **(Bottom-right)** Task distribution by disease category (upper) and task type (lower).

**Figure 3:**
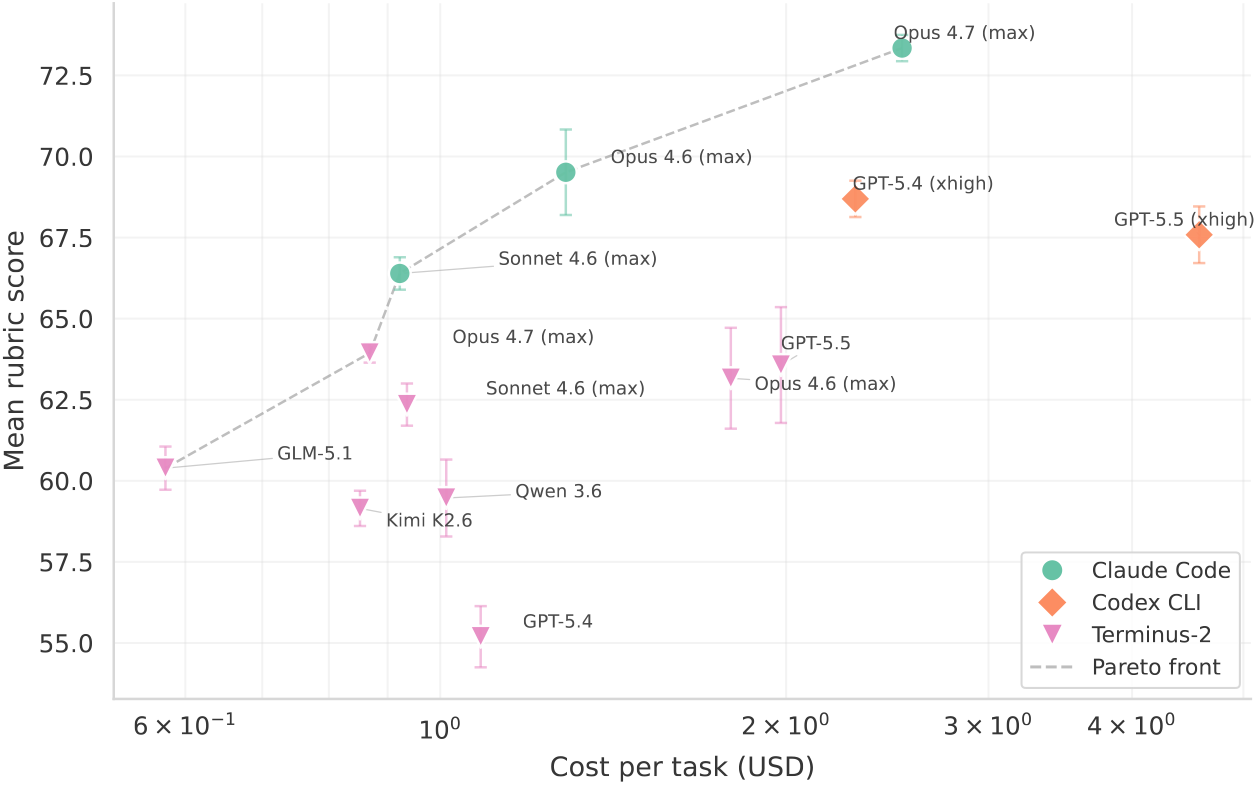
Cost vs. performance across agent–model configurations. Mean rubric score versus mean cost per task (USD, log-scale). Error bars show standard deviation across 3 reruns. Points above the dashed Pareto front dominate cheaper configurations. Gemini-based configurations are omitted because cost data was unavailable for the Gemini CLI runs.

#### Analyzing Biomni user traces

We analyzed 32,014 user queries from the open-source Biomni platform [Huang et al., 2025] to identify what scientists actually ask agents to do. Data analysis dominated this distribution, motivating BiomniBench-DA as the framework’s first release. Top biomedical domains include oncology, immunology, and neuroscience, and common tasks span differential expression and pathway analysis alongside a long tail of specialized analyses (see Appendix D).

#### Paper selection

Guided by these patterns, we selected source papers along four criteria. (1) The paper’s analysis matches one of the common task types observed in the Biomni user traces. (2) The selections collectively cover diverse disease domains and analytical methods, spanning both basic-science and translational-medicine venues so the benchmark is relevant to both academic and industry researchers. (3) The underlying data is publicly available through repositories such as GEO, ArrayExpress, or the paper’s supplementary materials. (4) The paper appears in a high-impact venue (Nature, Cell, Science, and their family journals such as Nature Medicine, Cancer Cell, Science Immunology) or as a recent preprint of similar caliber. The 100 tasks come from 21 such publications (full list in Appendix Table 4).

#### Expert curation

For each selected paper, we recruit either an original author of the paper or a domain expert with 5+ years of research experience. The expert is responsible for both formulating the research question and preparing the underlying dataset. We set expectations by sharing example questions and source papers up front, then ask the expert to read the paper carefully, locate the underlying dataset, and draw on both the paper’s analyses and their own domain expertise to generate the questions a researcher in the field would ask. The goal is not necessarily to reproduce the paper’s analyses but to capture a range of difficulty and topical diversity, including questions the paper did not directly address but the data could support.

#### Ground-truth generation

We ask the expert to produce a reference analytical trace as a Jupyter notebook or R Markdown document, documenting both the analysis code and the reasoning process throughout, alongside the final answer and a biomedical interpretation that extends beyond what the source paper reports. Each trace is verified to run on the public dataset and reviewed across multiple rounds by independent reviewers to ensure correctness.

#### Rubric design

Each task includes a task-specific rubric of 5–10 criteria, each anchored at a key decision point in the analysis and tagged to one or more of the six evaluation dimensions: data handling, method selection, statistical rigor, biological interpretation, scientific reasoning, and source reliability. Every criterion has three levels: **A** (a fully correct answer), **B** (right intention but with a minor mistake or partial match), and **C** (the step is skipped or completely wrong). Per-criterion point totals are set by the curating expert according to importance: A awards full credit, B usually half, and C zero. Each level enumerates multiple correct alternatives where applicable, so an agent that takes a sound but different analytical path is not penalized. Rubrics are finalized after multiple rounds of review and revision.

### 2.2 Evaluation protocol

#### Standardized agent execution

After collecting all task instructions and rubrics, we converted each task into the **Harbor** execution framework, which provides both the sandboxed runtime environment and the agent harnesses we evaluate (Codex CLI, Terminus-2, Claude Code, Gemini CLI). Each task is provisioned in a Docker container with the task description, the public dataset, and a single Markdown instruction file (an example appears in Appendix A). The container has Python 3 and R pre-installed, 2 CPUs, internet access, and a one-hour execution budget, sized so most tasks finish well within it. The task instruction itself prescribes the output format: each agent must write `trace.md` (the analytical narrative: decisions, code, intermediate findings, and interpretation) and `answer.txt` (a structured short-form response to the primary question) on completion, following the headings and detail level shown in the appendix example. In our pilot runs all of the LLM bases we tested produced well-formed trace and answer files, so the same output schema applies uniformly across harnesses. The instruction also explicitly forbids the agent from searching for or reading the source publication that the task was curated from, and spot-checks of agent traces showed no clear evidence of paper-derived shortcuts.

#### LLM judge

We use an LLM judge to score each agent trace against its task-specific rubric. For each criterion, the judge is given the rubric together with the agent’s full `trace.md` and `answer.txt`, and is asked to select the A/B/C level that best describes the agent’s work (prompt template in Appendix A). The judge returns only the categorical level. Per-criterion points and the final 0–100 task score are computed programmatically from the rubric’s level-to-points table. To choose the judge model, we compared five candidate LLMs (Claude Opus 4.6, DeepSeek V3.2, Gemini 3.1 Pro, GPT-5.4, Qwen 3.6) against internal-expert A/B/C ratings on 597 rubric criteria drawn from 35 tasks. Gemini 3.1 Pro showed the strongest agreement with humans on every metric (per-criterion exact accuracy 82%, linear-weighted Cohen’s *κ* = 0.70, quadratic-weighted *κ* = 0.73), exceeding the next-best candidate (Qwen 3.6) by 0.02 on accuracy, 0.06 on linear *κ*, and 0.04 on quadratic *κ*. We adopt Gemini 3.1 Pro as the judge throughout the paper. Full per-judge metrics and protocol details appear in Appendix B, Table 3.

### 2.3 Dataset statistics

BiomniBench-DA spans a broad cross-section of biomedical data analysis: 100 tasks distributed across 5 disease areas and a general-biology category, and 17 task types, drawn from 21 high-impact publications (Figure 2 bottom, Appendix Table 4).

#### Disease domains

The 100 tasks are distributed across five disease areas and a general-biology category for tasks not tied to a single disease: oncology (47%: colorectal cancer, NSCLC, melanoma, pan-cancer proteogenomics), metabolic and endocrine disease (17%: gender-affirming hormone therapy, glycemic-response phenotyping, NAFLD), immunology (14%: sepsis, autoimmune T-cell biology, SLE), general biology (10%: phase-separation predictor screening, drug-response landscape in primary human cells), neurology (8%: ALS), and cardiovascular (4%: multi-omic response to endurance training).

#### Analytical task types

The 100 tasks fall into 17 task types. The six most common cover 55% of the benchmark and represent the workhorse analyses that appear across data modalities in everyday biomedical research: association testing (11%), mutation analysis (10%), multi-omic integration (10%), pathway enrichment (8%), differential expression (8%), and clustering (8%). The remaining 45% is a long tail of more specialized analyses: predictive modeling, cell-composition analysis, survival analysis, chromatin profiling, signature scoring, longitudinal analysis, cross-cohort comparison, GWAS–eQTL colocalization, T-cell receptor repertoire analysis, co-expression network construction, and cell–cell communication inference.

## 3 Experiments and results

We evaluate BiomniBench-DA across nine frontier base models and four agent harnesses. Section 3.1 fixes the harness to isolate the effect of base-model capability, Section 3.2 varies model and harness jointly, and Section 3.3 decomposes performance by task type and evaluation dimension.

### 3.1 Performance of base LLMs with fixed harness

We first investigate the performance of different base LLMs by pairing them with a common codingagent harness, Terminus-2. Table 1 reports mean rubric scores for Terminus-2 paired with nine frontier base models, averaged across three independent reruns of the full 100-task benchmark.

**Table 1:** Terminus-2 agent benchmark on BiomniBench-DA across 9 frontier models. Score is the mean rubric score across 3 reruns of 100 tasks. *±* is the standard deviation across the three replicate means. Per-task SD is the mean over the 100 tasks of the within-task standard deviation across reruns. Time, cost, and turns are per-task medians. All runs use the default setting described in Appendix A.

| Model | Score (mean ± SD) | Per-task SD | Time/task | Cost/task | Turns |
| --- | --- | --- | --- | --- | --- |
| Claude Opus 4.7 | 63.94 ± 0.30 | 7.3 | 8.0m | \$0.87 | 15 |
| GPT-5.5 | 63.57 ± 1.79 | 9.4 | 11.4m | \$1.98 | 16 |
| Claude Opus 4.6 | 63.16 ± 1.56 | 8.3 | 17.0m | \$1.79 | 24 |
| Claude Sonnet 4.6 | 62.35 ± 0.65 | 7.5 | 13.6m | \$0.94 | 20 |
| GLM-5.1 | 60.39 ± 0.67 | 8.8 | 10.6m | \$0.58 | 20 |
| Qwen 3.6 | 59.47 ± 1.19 | 8.2 | 23.6m | \$1.01 | 22 |
| Kimi K2.6 | 59.15 ± 0.54 | 9.8 | 25.0m | \$0.85 | 31 |
| GPT-5.4 | 55.19 ± 0.94 | 11.5 | 12.6m | \$1.08 | 13 |
| Gemini 3.1 Pro | 44.27 ± 0.92 | 7.5 | 4.9m | \$0.37 | 12 |

Closed-source frontier models lead but are tightly clustered: Claude Opus 4.7 (63.94), GPT-5.5 (63.57), Claude Opus 4.6 (63.16), and Claude Sonnet 4.6 (62.35) sit within 1.6 points of each other. The strongest open-weight models (GLM-5.1 60.39, Qwen 3.6 59.47, Kimi K2.6 59.15) trail by only 3–5 points, at substantially lower cost (e.g., GLM-5.1 at $0.58/task vs. Opus 4.7 at $0.87/task). Still, even the best configuration scores below 64 of 100, leaving substantial headroom across the benchmark. Gemini 3.1 Pro is an outlier: it produces the shortest traces (median 4.9 minutes, 12 turns) but lands at 44.27, well below the other models. Per-task SDs range from 7.3 to 11.5 points, indicating substantial run-to-run variability. Newer model generations are also more efficient at similar score levels: Opus 4.7 reaches 63.94 in a median 15 turns and 8.0 minutes, while Opus 4.6 reaches a comparable 63.16 in 24 turns and 17.0 minutes.

### 3.2 Performance across base LLMs and agent harnesses

We next examine the joint effect of base LLM and agent harness by pairing each closed-source frontier coding agent (Claude Code, Codex CLI, Gemini CLI) with its strongest base LLMs, all at maximum reasoning effort (Appendix A).

The top configuration is Claude Code paired with Opus 4.7 at 73.34, followed by Claude Code with Opus 4.6 (69.51), Codex CLI with GPT-5.4 (68.69), Codex CLI with GPT-5.5 (67.59), and Claude Code with Sonnet 4.6 (66.39). Holding the base model fixed, swapping the harness moves scores substantially. The clearest case is GPT-5.4: 68.69 under Codex CLI versus 55.19 under Terminus-2, a 13.5-point spread attributable entirely to agent architecture. For Claude Opus 4.7, the Claude Code–Terminus-2 gap is 9.4 points (73.34 vs. 63.94). These per-model harness gaps exceed the gap between successive Anthropic model generations under the same harness (Opus 4.7 vs. Opus 4.6 under Claude Code: 3.8 points), establishing the agent harness as a primary design choice rather than a thin wrapper around the base model. Switching to these harnesses also reshapes the runtime profile: Codex CLI runs use far more turns than Terminus-2 on the same model. For example, GPT-5.4 jumps from 13 turns under Terminus-2 to 82 under Codex CLI, and GPT-5.5 from 16 to 53, with cost rising in step (GPT-5.5 from $1.98 to $4.58 per task). Claude Code is closer to Terminus-2 in turn count but still trends upward. For Opus 4.7, turns rise from 15 to 34 and cost from $0.87 to $2.52 per task, and its highest-scoring configurations are also the most expensive.

Codex CLI provides the largest boost over Terminus-2 on its native models. GPT-5.4 gains 13.5 points (from 55.19 to 68.69) while GPT-5.5 gains only 4.0 (from 63.57 to 67.59), reversing the ordering seen under Terminus-2 (where GPT-5.5 outperformed GPT-5.4) and suggesting that a well-tuned harness can partly compensate for base-model weaknesses.

### 3.3 Anatomy of agent performance

To see where agents succeed and fall short, we decompose performance by task type (Figure 4a) and by evaluation dimension (Figure 4b).

**Figure 4:**
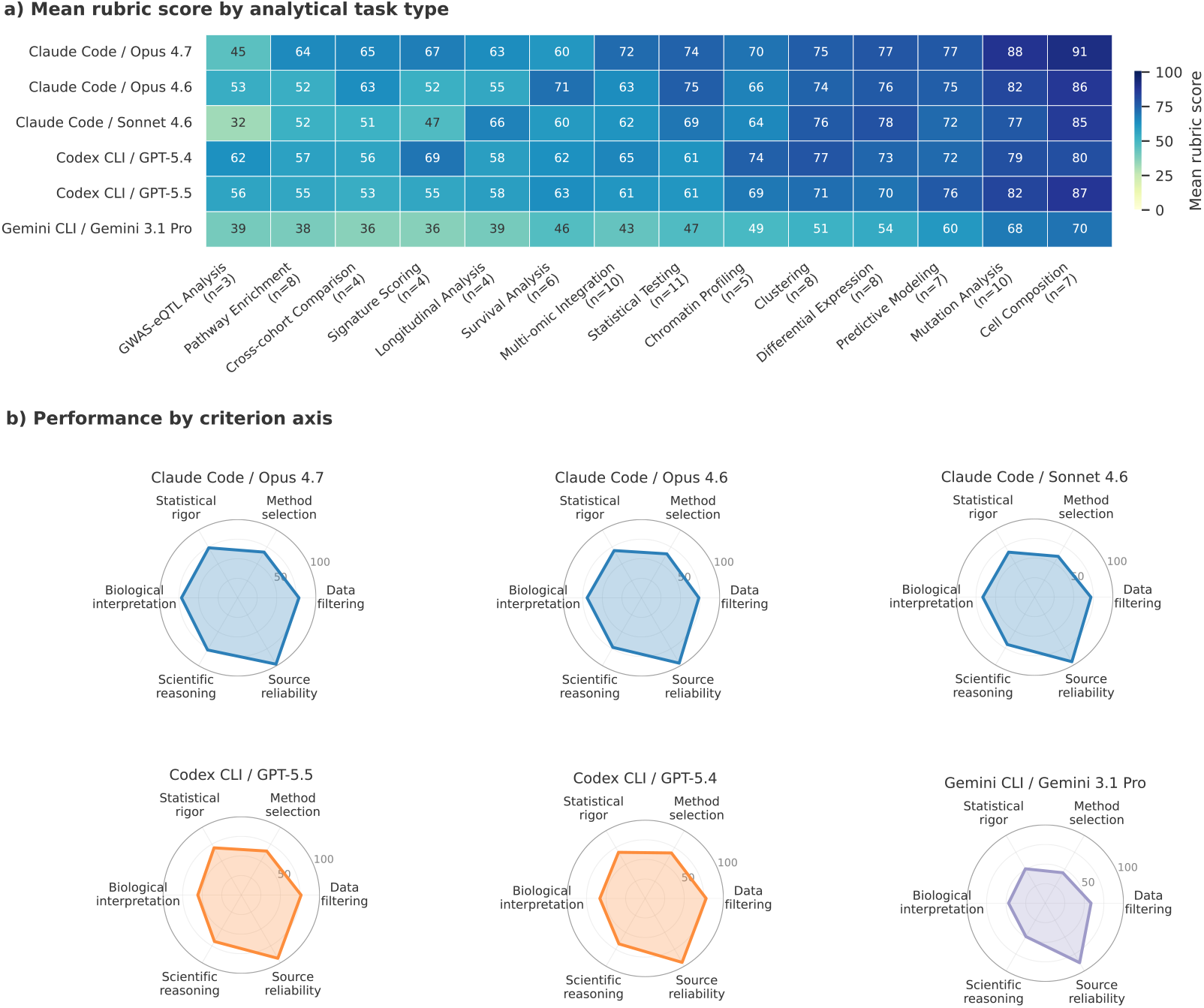
Anatomy of agent performance across cross-harness configurations. **(a)** Mean rubric score by task type for the six configurations (categories with at least 3 tasks, sorted hardest to easiest). **(b)** Percentage of available rubric points earned along the six evaluation dimensions (data handling, method selection, statistical rigor, biological interpretation, scientific reasoning, source reliability).

Performance varies substantially across task types. Two categories are universally hard: GWAS– eQTL Analysis (best score 65 across all six configurations) and Pathway Enrichment (best 67). Even the strongest agents drop at least 10 points on these relative to their overall mean. Two are nearly maxed out: Cell Composition (max 91, min 70) and Mutation Analysis (max 86, with the top five configurations all at or above 80). The largest harness gap appears on Cross-cohort Comparison (Claude Code spread 32–53 vs. Codex CLI 51–56), suggesting that agent harnesses vary most on tasks requiring orchestration across multiple datasets.

By evaluation dimension, a consistent picture emerges across agents: they reliably ground their analyses in identifiable sources and handle diverse data formats well, but consistently fall short on selecting the appropriate analytical method, applying scientific reasoning, and interpreting results in biological context. The numbers reflect this: *source reliability* earns 88–98% of available points across all configurations and *data handling* earns 58–78%, while *method selection* sits at the bottom (44–67%) and statistical rigor, biological interpretation, and scientific reasoning occupy a middle band (45–74%). Claude Code and Codex CLI differ only at the margins, with Claude Code stronger on biological interpretation and Codex CLI matching Claude Code on data handling and method selection. A qualitative review of the lowest-scoring criteria across configurations distills these deficits into three recurring failure patterns: wrong method selection, flawed biological interpretation, and poor scientific reasoning, with concrete cases in Appendix C.

## 4 Discussion

Current frontier LLM agents show progress on biomedical data analysis: the strongest closed-source and open-weight base models cluster within a few points of each other under a fixed harness, complete multi-step analyses end-to-end, and reliably ground their claims in identifiable sources. The gap to expert-level performance, however, remains substantial. Even the best configuration scores below 75 of 100 on average, with consistent weak spots in choosing the right analytical method, interpreting results in biological context, and applying scientific reasoning across multi-step procedures.

Closing this gap is not only about scaling the base model: the agent harness around it materially shapes how reliably the analysis is completed. The 13.5-point Codex CLI / Terminus-2 gap for GPT-5.4 exceeds the 3.8-point Opus 4.7 / Opus 4.6 generational gap under Claude Code, and execution failures (data handling, tool misuse) shrink with structured scaffolding while reasoning failures persist regardless of harness. The general-purpose coding harnesses we tested were not designed for biomedical research, and there is substantial headroom for a biomedical-aware harness that builds in domain conventions, including specialized data-loading utilities, statistical-default selection, and structured biological-interpretation steps.

BiomniBench is modular, and future work will broaden the framework along several axes. First, we will release additional task types beyond data analysis: experimental design, literature synthesis, and protocol optimization tasks such as protein design and assay development. Second, the dimension-level diagnostics point to concrete improvement targets for both base models and harnesses on the weak axes (method selection, biological interpretation, scientific reasoning). Third, we will extend the benchmark to more diverse, longer-horizon, and compute-intensive tasks that require multi-turn collaboration with the user rather than a single one-shot attempt, mirroring how researchers iterate with an agent in practice.

Several limitations remain. Rubric construction is a considerable expert effort, and it is not always feasible to enumerate every defensible analytical path. Alternative correct approaches may exist beyond those the rubric explicitly credits. The LLM judge agrees strongly with human experts overall (Cohen’s *κ* = 0.70, Appendix B) but is not a substitute for expert judgment on the most subjective dimensions. For high-stakes evaluation we recommend supplementing automated scoring with human review. Finally, expert curation, rubric authoring, and reviewer rounds remain labor-intensive, and the per-task cost of producing a benchmark item at this depth is substantial. A natural next step is to use agents themselves to assist with benchmark construction, drafting candidate tasks and rubrics from the source literature for expert review, so that the framework can scale to broader task types and larger task pools.

## 5 Related work

Knowledge benchmarks such as HLE [Phan et al., 2025], GPQA [Rein et al., 2023], Frontier-Science [Wang et al., 2026], PubMedQA [Jin et al., 2019], and BioASQ [Tsatsaronis et al., 2015] test factual recall through multiple-choice (MCQ) formats and do not evaluate end-to-end research execution. LAB-Bench and its successor [Laurent et al., 2024, 2026] extend coverage to practical research skills (literature retrieval, figure interpretation, protocol design) but remain mostly MCQ-based.

A newer generation evaluates multi-step analytical tasks: BixBench [Mitchener et al., 2025], BioML-Bench [Miller et al., 2025], BioAgent Bench [Fa et al., 2026], scBench [Workman et al., 2026], and SpatialBench [Workman et al., 2025]. All score the final answer, which cannot distinguish a correct process from an outcome-only shortcut, and benchmark ground truth itself is often noisy or contested, compounding the problem [Tu et al., 2026]. Concurrent work explores adjacent directions. GeneBench introduces intermediate decision points along the analysis trajectory in genomics and quantitative biology, but grades each step by exact match against a single reference answer [Li and Ho, 2026]. CompBioBench targets real-world computational-biology problems but operates on synthetic data and likewise scores outcomes by exact match [Nair et al., 2026]. BioMysteryBench probes biomedical-agent performance through investigative bioinformatics scenarios [Anthropic, 2026]. BiomniBench is complementary: it grades the full analytical trajectory on real-world data through expert-authored, ordinal A/B/C rubrics that explicitly credit alternative valid analytical paths.

## 6 Conclusion

We introduced **BiomniBench**, a process-level evaluation framework for assessing LLM agents’ capability on real-world biomedical research, and released it as **BiomniBench-DA**: 100 data analysis tasks curated from papers in Nature, Cell, Science, and similar journals, co-developed with the original authors and domain experts. Across nine frontier base models and four agent harnesses, closed-source and open-weight bases cluster within a few points of each other, and the agent harness can shift scores by more than the gap between successive model generations. Even the best configuration scores below 75/100 on average, with consistent weak spots on *method selection, biological interpretation*, and *scientific reasoning* despite reliable source grounding. By grading the full analytical trajectory and crediting alternative valid paths rather than a single reference answer, BiomniBench moves evaluation beyond outcome-only leaderboards and yields the dimension-level diagnostics needed to direct future base-model and harness development toward the capabilities biomedical research demands.

## A Experimental setup and examples

### A.1 Run protocol

#### Harbor framework

All agent runs are executed within Harbor v3, a sandboxed execution framework. Each task is provisioned in an isolated Linux container with the task description, the public dataset, and task-specific runtime constraints (time, cost, and turn budgets). Every agent is instructed to write `trace.md` (the analytical narrative: decisions, code, intermediate findings, interpretation) and `answer.txt` (a structured short-form response to the primary question) on completion.

#### Effort settings

Anthropic models in Terminus-2 (Table 1) use the `terminus2_adaptive` variant with adaptive interleaved thinking and `effort=max`. Other Terminus-2 runs use stock Terminus-2 with `reasoning_effort=high`, the harness’s maximum supported setting (Gemini maps this to `thinkingBudget=24576`, and for OpenRouter open-weight models it is forwarded to the provider). For cross-harness runs (Table 2), Claude Code uses `effort=max`, Codex CLI uses `reasoning_effort=xhigh`, and Gemini CLI uses `thinkingBudget=-1`.

**Table 2:** Cross-harness benchmark on BiomniBench-DA. Six agent–model configurations across Claude Code, Codex CLI, and Gemini CLI. Conventions match Table 1. Run details in Appendix A.

| Harness | Model | Score (mean ± SD) | Per-task SD | Time/task | Cost/task | Turns |
| --- | --- | --- | --- | --- | --- | --- |
| Claude Code | Opus 4.7 | 73.34 ± 0.41 | 8.6 | 10.2m | \$2.52 | 34 |
| Claude Code | Opus 4.6 | 69.51 ± 1.32 | 7.9 | 8.3m | \$1.29 | 17 |
| Codex CLI | GPT-5.4 | 68.69 ± 0.56 | 8.6 | 14.3m | \$2.30 | 82 |
| Codex CLI | GPT-5.5 | 67.59 ± 0.87 | 8.7 | 8.8m | \$4.58 | 53 |
| Claude Code | Sonnet 4.6 | 66.39 ± 0.50 | 7.8 | 8.4m | \$0.92 | 23 |
| Gemini CLI | Gemini 3.1 Pro | 49.83 ± 1.33 | 8.9 | 5.4m | – | – |

#### Accuracy calculation

Each agent–model configuration is run three times independently on the full 100-task benchmark. The headline score (the “Score” column in Tables 1–2) is computed in two steps: for each task we average rubric scores across the three reruns, then average those per-task means over all 100 tasks. The *±* value is the standard deviation across the three full-benchmark replicate means (one per rerun), capturing rerun-to-rerun stability. The per-task SD column is the mean over the 100 tasks of the within-task standard deviation across reruns, capturing typical within-task variability.

#### Rerun on transient failure

Tasks that fail to produce a valid `trace.md / answer.txt` pair (container timeout, agent crash, or invalid output format) are automatically rerun. If the retry also fails, the task is recorded as a hard failure for that rep and contributes a zero rubric score.

#### Time and cost calculation

Per-task execution time, cost, and turn count are reported as medians: for each task we take the median across the three reruns, then the median over the 100 tasks. Cost is measured from API token usage. Gemini CLI does not expose token usage and is reported without cost data.

### A.2 Example task instruction (Harbor format)

Each BiomniBench-DA task is delivered to the agent as a single `instruction.md` file containing the research question, the data-file manifest, and the required output format (the prompt that asks the agent to produce `trace.md and answer.txt)`. We reproduce the full instruction for da-1-1 (CRC anti-PD-1, *Cell* 2024; GSE236581) verbatim:

~~~
# Task: Spatiotemporal Single-Cell Analysis of Immunotherapy Response in CRC
## Question
At baseline (pre-treatment), are there significant differences in the proportion of T cells among all immune cells in tumor tissues between patients of different response groups? What are the changing trends of different response groups before and after treatment?
## Data Files
Single-cell cohort from the spatiotemporal CRC anti-PD-1 study
(Cell 2024; GSE236581): pre- and on-treatment tumor, adjacent normal,
and peripheral blood samples.
 -‘GSE236581_counts.mtx’: Raw UMI counts in Matrix Market sparse format (genes x cells).
 -‘GSE236581_barcodes.tsv’: Cell barcodes (one per line, ordered to match columns of the counts matrix). Format ‘{SampleID}_{CellBarcode}’.
 -‘GSE236581_features.tsv’: Gene index with columns ‘gene_id’, ‘gene_symbol’, ‘feature_type’; ordered to match rows of the counts matrix (no header row).
 -‘GSE236581_CRC-ICB_metadata.txt’: Space-separated cell-level metadata. Columns: ‘orig.ident’, ‘nCount_RNA’, ‘nFeature_RNA’, ‘Ident’, ‘Patient’, ‘Treatment’ (I = pre-treatment, II-IV = on-treatment), ‘Tissue’ (Tumor / Normal / Blood), ‘MajorCellType’, ‘SubCellType’.
 -‘clinical.xlsx’: Patient and sample metadata. Three sheets (header is row 2):
   -‘scRNA-seq patient meta’: clinical variables per patient (Response CR/PR/SD/PD, Tumor Regression Ratio, TMB, MSI/MSS, Treatment Regimen).
   -‘scRNA-seq sample meta’: per-sample info (Biopsy Site, Treatment Stage, Treatment point, Sampling approach).
   -‘Validation patient meta’: independent validation cohort with Response and Sample Type.
## Required Outputs
You MUST create the following two output files:
### 1. Analysis Trace (‘/app/trace.md’)
Document your complete analysis process in markdown format, following this structure. This trace is evaluated by an expert judge: thoroughness and clarity directly affect your score.
Required sections (use these as headings in your trace):
1. Objective: Restate the question; define what “success” looks like.
2. Data Sources: For each file: name, dimensions, key columns, example values for any column you’ll filter or group by, data quality notes.
3. Approach: Numbered analytical steps. For each step, document:
   -Description: what was done.
   -Decision and rationale: which method, threshold, normalization, or filter was chosen, and why; what alternatives were considered or rejected; any assumption or judgment call.
   -Code: paste the actual code (Python / R / shell) used for each non-trivial operation in this step (filters, aggregations, joins, statistical tests, transformations, plotting calls). Use real, copy-pasteable code (not pseudocode and not a paraphrase).
   -Quantitative intermediate result (counts, p-values, dimensions after filtering).
4. Results: Specific quantitative findings (numbers, gene/protein names, statistics, summary tables); biological / clinical interpretation; what the analysis cannot conclude (limitations).
5. References: Real, identifiable citations (author + year, DOI, PubMed ID, named database) for biological mechanisms invoked, methodology choices, and any external knowledge used.
Key principles:
-Show intermediate result counts at every filtering step
(e.g., “1,850 -> 1,720 -> 1,680”).
-Name specific entities and quantities; avoid vague summaries.
-Justify analytical choices (test selection, threshold, normalization).
-Ground biological / clinical claims in real references; do not assert from memory.
-State what the analysis cannot conclude (limitations).
### 2. Final Answer (‘/app/answer.txt’)
Write your final answer to the question above in plain text format.
## Environment
-Python 3 and R are pre-installed.
-Install any additional packages you need (pandas, statsmodels, etc.).
-Internet access is available for package installation.
-Do not search for or read the specific source paper, figures, or supplementary materials that the dataset comes from. Solve the task directly from the provided data and domain knowledge.
~~~

### A.3 Example rubric (full)

The full rubric for da-1-1 contains nine criteria summing to 100 points (Criteria 1–8 carry positive weight; Criterion 9 is a non-positive “Source reliability” check that can deduct points). Each criterion has three levels [A]/[B]/[C] with rubric-defined point values. Across the benchmark, rubrics contain 5–10 criteria.

~~~
RUBRIC: T Cell Proportion Analysis in Tumor Tissue Across PD-1 Blockade
        Response Groups
Total Points: 100/100
CRITERIA (9):
Criterion 1: Clinical Data Integration with Cell-Level Metadata
  Description: Evaluates correct merging of patient-level clinical data
  (clinical.xlsx containing Response, TRG status) with cell-level metadata,
  preserving the cell ID as a usable key after the merge.
  Levels: A=11 B=7 C=0
    [A]: Correctly merges single-cell metadata with clinical data on the Patient field, ensures the cell identifier remains accessible (as index or column) after merging, and cleans irrelevant columns.
    Any pandas / R / numpy idiom is acceptable.
    [B]: Merges clinical data but with errors in cell-identifier preservation, incomplete column cleaning, or partially incorrect merge key.
    [C]: Does not integrate clinical data or integration is fundamentally incorrect, preventing downstream analysis.
Criterion 2: Baseline Subset Selection
  Description: Evaluates correct identification and filtering of cells for baseline analysis: restricting to Tumor tissue, Baseline timepoint, and immune cell lineages.
Levels: A=16 B=10 C=0
   [A]: Correctly applies all three filters: Tissue == ‘Tumor’,
Treatment == ‘I’ (Baseline), and MajorCellType restricted to immune lineages (T, B, Mye, ILC). Demonstrates understanding that ‘I’ represents the baseline/pre-treatment timepoint.
   [B]: Applies some but not all required filters, or uses incorrect filter values (e.g., wrong Treatment stage for baseline, missing one immune cell type).
   [C]: Fails to filter data appropriately, or uses the entire dataset without subsetting for baseline tumor immune cells.
Criterion 3: Patient-Level Proportion Calculation
  Description: Evaluates the critical step of aggregating single-cell
data to patient-level proportions, correctly computing T cell
proportion as T_count / Total_Immune rather than using raw cell counts.
Levels: A=19 B=12 C=0
    [A]: Aggregates data to the patient level (not cell level), correctly defines the denominator as the sum of all immune-cell-lineage counts per patient (Total_Immune = T + B + Mye + ILC), and computes T_Cell_Proportion = T_count / Total_Immune per patient.
    [B]: Aggregates to patient level but makes errors in denominator calculation (e.g., includes non-immune cells), or computes raw counts instead of proportions.
    [C]: Uses cell-level data directly without patient-level aggregation, or fundamentally miscalculates the proportion metric.
Criterion 4: Appropriate Statistical Test Selection
  Description: Evaluates the choice and justification of statistical test for comparing T cell proportions across response groups at baseline, considering small sample sizes and non-normal distributions.
Levels: A=13 B=8 C=0
    [A]: Uses Mann-Whitney U test (stats.mannwhitneyu) or equivalent non-parametric test, with clear justification that the test is appropriate for small sample sizes and potentially non-normal data distributions.
    [B]: Uses a reasonable statistical test but without justification, or uses a parametric test (e.g., t-test) without verifying normality assumptions.
    [C]: No statistical testing performed, or uses a clearly inappropriate test.
Criterion 5: Pairwise Statistical Comparisons and Reporting
  Description: Evaluates execution of all required pairwise comparisons
and proper reporting of results.
Levels: A=14 B=9 C=0
    [A]: Defines and tests all three pairwise comparisons (CR vs PR, CR vs SD, PR vs SD); reports raw per-comparison p-values matching expectations (CR vs PR p=1.0 ns, CR vs SD p∼0.048, PR vs SD p∼0.033); applies a standard significance scheme (***: p<0.001, **: p<0.01, *: p<0.05, ns). Additionally reporting a multiple-testing-corrected p-value (Bonferroni or BH-FDR) alongside the raw values is acceptable.
    [B]: Tests some but not all pairwise comparisons, OR reports only adjusted p-values without showing the raw per-comparison values, OR reports p-values without any significance classification.
    [C]: Does not perform pairwise comparisons, or reports incorrect results.
Criterion 6: Longitudinal Data Preparation and Temporal Ordering
  Description: Evaluates the preparation of data for longitudinal
analysis, including re-filtering for all timepoints, recalculating
proportions, and enforcing logical temporal order.
Levels: A=13 B=8 C=0
    [A]: Re-filters data to include all four treatment timepoints (I, II, III, IV) for tumor tissue immune cells; re-computes T cell proportions per patient per timepoint using the same aggregation method as at baseline; enforces a logical temporal order (I -> II -> III -> IV) when reporting and plotting.
    [B]: Includes multiple timepoints but misses some, or does not enforce temporal ordering, or reuses baseline-only filtered data.
    [C]: Does not perform longitudinal analysis, or uses only baseline data.
Criterion 7: Interpretation of Baseline Findings
   Description: Evaluates the biological interpretation of baseline statistical results in the context of anti-PD-1 immunotherapy.
Levels: A=7 B=4 C=0
    [A]: Correctly interprets that at baseline, CR and PR groups show no significant difference in T cell proportions (p=1.0), while both CR and PR groups differ significantly from SD (p<0.05). Provides biological interpretation linking higher baseline T cell infiltration in responders (CR/PR) to a “hotter” tumor immune microenvironment, and discusses implications for predicting treatment efficacy.
   [B]: Correctly reports which comparisons are significant vs. not significant, but provides limited or no biological interpretation.
   [C]: Misinterprets statistical results or provides no interpretation of baseline findings.
Criterion 8: Interpretation of Longitudinal Trends
  Description: Evaluates the characterization and interpretation of temporal dynamics in T cell infiltration across response groups following PD-1 blockade therapy.
Levels: A=7 B=4 C=0
    [A]: Characterizes distinct temporal patterns for each response group (e.g., how T cell proportions evolve from Stage I through Stage IV for CR, PR, and SD), identifies differences in T cell recruitment and expansion dynamics between response types, and discusses the association between temporal evolution of T cell abundance and clinical treatment responses.
    [B]: Describes general trends but without group-specific characterization or without linking to clinical response mechanisms.
    [C]: No interpretation of longitudinal trends, or interpretation contradicts the data.
Criterion 9: Source Reliability
  Description: Evaluates whether the agent grounds its values,
annotations, and biological / clinical interpretations in identifiable
sources, the provided data files, named public databases, or real peer-reviewed references, rather than asserting facts from memory without source.
Levels: A=0 B=-5 C=-10
    [A]: Numerical results (cell counts, proportions, p-values) are traceable to the provided data (clinical.xlsx, GSE236581 metadata, count matrix) or a documented transformation.
Cell-type labels and treatment-stage codes come from the provided metadata, not assumed. Biological and clinical interpretations are supported by real, identifiable references (author + year, DOI, PubMed ID, or named review article).
    [B]: Most numerical results and labels are traceable, but some interpretive claims are asserted without source attribution or cite only vague phrases like “studies show”.
    [C]: Reports values, cell-type labels, or biological claims with no clear source. Hallucinated content (made-up labels, invented p-values, or fabricated citations) mixed with computed results.
~~~

### A.4 Example judge prompt and output

Each task is scored by the LLM judge using the following prompt template (the rubric, the agent’s `trace.md`, and the agent’s `answer.txt` are spliced in at runtime). The judge returns a JSON object that selects one level per criterion together with a one-sentence justification; per-criterion points are then computed deterministically from the level letters using the rubric’s point table.

~~~
You are an expert evaluator for a data analysis task.
Evaluate the agent’s work using the following rubric:
{rubric}
Here is the agent’s analysis trace:
<trace>
{trace_content}
</trace>
Here is the agent’s final answer:
<answer>
{answer_content}
</answer>
For each criterion in the rubric, choose ONE level: A, B, or C - based purely on which level description best describes the agent’s work. Do not output numerical points; the score for each level is computed automatically from the rubric.
You MUST respond with a JSON object in exactly this format:
{
“criteria”: {
         “criterion_1”: {“level”: “A”, “reason”: “<one-sentence explanation>“},
         “criterion_2”: {“level”: “B”, “reason”: “<one-sentence explanation>“},
         …
   },
   “overall_reasoning”: “<short summary>“
}
Each “level” value must be exactly the single character “A”, “B”, or “C”. Only output the JSON object, nothing else.
~~~

A representative judge response on da-1-1:

~~~
{
  “criteria”: {
          “criterion_1”: {“level”: “A”, “reason”: “Merges single-cell metadata and clinical.xlsx on Patient; cell IDs preserved.”},
          “criterion_2”: {“level”: “A”, “reason”: “Filters Tissue=Tumor, Treatment=I, MajorCellType in {T,B,Mye,ILC}.”},
          “criterion_3”: {“level”: “A”, “reason”: “Patient-level T/Total_Immune with denominator = T+B+Mye+ILC.”},
          “criterion_4”: {“level”: “A”, “reason”: “Mann-Whitney U with justification for non-normal small samples.”},
          “criterion_5”: {“level”: “B”, “reason”: “Reports BH-FDR but omits raw per-comparison p-values.”},
          “criterion_6”: {“level”: “A”, “reason”: “Re-filters all four timepoints (I-IV); preserves I->II->III->IV order.”},
          “criterion_7”: {“level”: “A”, “reason”: “Interprets CR/PR vs SD baseline difference; links to hot-tumor framework.”},
          “criterion_8”: {“level”: “B”, “reason”: “Describes general temporal trends but no group-specific dynamics.”},
          “criterion_9”: {“level”: “A”, “reason”: “Numerical results traceable to provided files; references checkable.”}
      },
      “overall_reasoning”: “Strong baseline analysis with correct integration,
                             filtering, and statistical testing. Criterion 5
                             missed because raw per-comparison p-values are
                             omitted in favor of BH-corrected ones; Criterion 8
                             describes trends but not group-specific dynamics.”
  }
~~~

Per the rubric’s point table, this assignment yields a raw task score of 11 + 16 + 19 + 13 + 9 + 13 + 7 + 4 + 0 = 92*/*100, which is the per-task score reported in Tables 1–2. The same per-criterion points are also aggregated by dimension tag (each criterion is tagged to one or more of the six dimensions defined in Section 2.1) to produce the dimension-level scores shown in Figure 4.

## B LLM-judge validation

### Setup

We assembled a held-out evaluation set of 597 rubric criteria spanning 35 tasks. For each task, the internal domain experts who had originally constructed the task and its rubric were given the agent’s `trace.md` and `answer.txt` and instructed to judge every criterion on the A/B/C scale defined in Section 2.1. The same set was then independently rated by five candidate LLM judges: Claude Opus 4.6, DeepSeek V3.2, Gemini 3.1 Pro, GPT-5.4, and Qwen 3.6. Each judge was prompted with the task-specific rubric and the agent’s `trace.md` and `answer.txt`, and asked to return one A/B/C label per criterion with a one-sentence justification, using the prompt template reproduced in Appendix A.

### Metrics

We report two agreement metrics with the human gold standard: *Exact-match accuracy* at the criterion level (fraction of criteria where the judge’s label equals the human’s), and *Cohen’s κ* with linear and quadratic weights (see table caption for the weighting convention).

### Results

Gemini 3.1 Pro is the best candidate on every metric, with the gap to the runner-up widening on the ordinal-weighted *κ* scores (Table 3). Qwen 3.6 is competitive on raw exact accuracy but its overall *κ* drops because of a lenient bias toward “A”; GPT-5.4 and Claude Opus 4.6 occupy a middle band (*κ≈* 0.47–0.55); DeepSeek V3.2 is the weakest, with bimodal label distributions that essentially collapse the “B” class. We therefore use Gemini 3.1 Pro as the production LLM judge for all results in this paper.

**Table 3:** Judge agreement with human A/B/C ratings. (*n* = 597 criteria across 35 tasks). Cohen’s *κ* uses linear and quadratic weights, treating A/B/C as ordinal so an A*↔*C error is penalized more than an A*↔*B error.

| Judge | Exact acc. | Linear $\kappa$ | Quadratic $\kappa$ |
| --- | --- | --- | --- |
| **Gemini 3.1 Pro** | **0.82** | **0.70** | **0.73** |
| Qwen 3.6 | 0.80 | 0.64 | 0.69 |
| GPT-5.4 | 0.69 | 0.55 | 0.62 |
| Claude Opus 4.6 | 0.70 | 0.47 | 0.51 |
| DeepSeek V3.2 | 0.60 | 0.35 | 0.40 |

**Table 4:** BiomniBench-DA source publications. “Tasks” is the number of benchmark tasks derived from each paper.

| ID | Paper Title | Journal | Tasks |
| --- | --- | --- | --- |
| DA-1 | Spatiotemporal single-cell analysis decodes cellular dynamics underlying different responses to immunotherapy in colorectal cancer | Cancer Cell | 3 |
| DA-3 | Genomic and transcriptomic features of response to anti-PD-1 therapy in metastatic melanoma (Hugo et al.) | Cell | 4 |
| DA-4 | A single-cell atlas reveals immune heterogeneity in anti-PD-1-treated non-small cell lung cancer | Cell | 7 |
| DA-5 | Pan-cancer proteogenomics expands the landscape of therapeutic targets | Cell | 4 |
| DA-6 | Temporal dynamics of the multi-omic response to endurance exercise training | Nature | 4 |
| DA-8 | Individual variations in glycemic responses to carbohydrates and underlying metabolic physiology | Nature Medicine | 5 |
| DA-9 | Sotigalimab and/or nivolumab with chemotherapy in first-line metastatic pancreatic cancer (PRINCE trial) | Nature Medicine | 5 |
| DA-10 | Screening membraneless organelle participants with machine-learning models that integrate multimodal features | PNAS | 4 |
| DA-11 | KIR+CD8+ T cells suppress pathogenic T cells and are active in autoimmune diseases and COVID-19 | Science | 1 |
| DA-12 | Integrative analyses of noncoding RNAs reveal potential mechanisms augmenting tumor malignancy in lung adenocarcinoma | Nucleic Acids Res. | 4 |
| DA-13 | Plasma proteome adaptations during feminizing gender-affirming hormone therapy | Nature Medicine | 8 |

**Table 4 – continued from previous page**
| ID | Paper Title | Journal | Tasks |
| --- | --- | --- | --- |
| DA-14 | A consensus immune dysregulation framework for sepsis and critical illnesses | Nature Medicine | 7 |
| DA-15 | Integrative transcriptomic analysis of the amyotrophic lateral sclerosis spinal cord implicates glial activation and suggests new risk genes | Nat. Neurosci. | 8 |
| DA-16 | Transcriptomic profiling across the nonalcoholic fatty liver disease spectrum reveals gene signatures for steatohepatitis and fibrosis | Sci. Transl. Med. | 2 |
| DA-17 | Single-cell RNA-seq reveals cell type-specific molecular and genetic associations to lupus | Science | 6 |
| DA-18 | The Genomic Landscape of Endocrine-Resistant Advanced Breast Cancers | Cancer Cell | 7 |
| DA-19 | CBFβ-SMMHC Inhibition Triggers Apoptosis by Disrupting MYC Chromatin Dynamics in Acute Myeloid Leukemia | Cell | 8 |
| DA-20 | Mapping the Transcriptional Landscape of Drug Responses in Primary Human Cells Using High-Throughput DRUG-seq | bioRxiv | 6 |
| DA-24 | A genome-wide association study of neonatal metabolites | Cell Genomics | 2 |
| DA-25 | Dynamic prostate cancer transcriptome analysis delineates the trajectory to disease progression | Nat. Commun. | 2 |
| DA-26 | Building a translational cancer dependency map for The Cancer Genome Atlas | Nature Cancer | 3 |

### Failure modes

Beyond the headline metrics, the five judges fail in distinct ways. Claude Opus 4.6 collapses “B” into “A” (39% of human-B rated A); Qwen 3.6 is similarly lenient (43% of human-B rated A) despite its higher overall accuracy; GPT-5.4 is the inverse, harsh (24% of human-A rated B); DeepSeek V3.2 essentially eliminates the “B” class (21% B-recall, with 39% of human-B pushed all the way to C). Only Gemini 3.1 Pro produces a balanced confusion matrix, with diagonal recalls 87%*/*76%*/*73% and no off-diagonal cell exceeding 20%.

## C Failure mode case studies

The aggregate scores in Section 3 show *which* evaluation dimensions agents lose points on (Figure 4). This appendix documents *what those failures look like in practice* via concrete cases drawn from the benchmark.

### C.1 Wrong method selection

In the SUBSPACE immune-dysregulation study (DA-14-2), agents must determine whether immune dysregulation escalates monotonically with infection severity across an ordered four-level grade (healthy, inflammopathic, coagulopathic, adaptive). The question is specifically about monotonic ordered trend. Agents instead reach for Kruskal–Wallis or Mann–Whitney, then claim a robust monotonic relationship from a strongly significant omnibus p-value. The substitution is not a defensible alternative: Kruskal–Wallis tests whether *any* group differs, not whether differences increase monotonically with the ordered factor, so a significant K–W result is not evidence for the monotonic claim the agent makes. The Jonckheere–Terpstra trend test, run on the same data, returns a non-significant *p≈* 0.40, revealing within-group heterogeneity that the omnibus alternatives by construction cannot detect.

### C.2 Flawed biological interpretation

In the GAHT proteome study (DA-13-8), agents must determine whether protein changes induced by feminizing gender-affirming hormone therapy converge with or diverge from disease-associated proteomic signatures, anchored to specific clinical phenotypes (e.g., autoimmune disease, atherosclerosis, allergic asthma) and identifying the contributing proteins for each. Agents instead produce a proteome-wide convergence/divergence summary that does not anchor the analysis to the named phenotypes. This gap is not a difference in style: the question explicitly names the disease contexts of interest, and an answer at the proteome-wide level cannot speak to whether GAHT pushes the proteome toward or away from any one of those specific disease signatures, which is the clinically meaningful claim.

### C.3 Poor scientific reasoning

In the BRCA dependency-mapping task (DA-26-2), agents must identify biomarkers that are statistically significant in BRCA patient-level DepMap predictions but not in cell-line models. The task specifies a multi-step reasoning chain: a normality likelihood-ratio test on the dependency-score distributions, a classification step that decides whether each gene’s distribution is normal or t-skewed, a sum-of-squared-deviations computation that weights genes by their distributional shape, and integration with an external druggable-gene list. Agents replace this chain with a single Pearson or Spearman correlation between patient and cell-line scores. The substitution does not address the question being asked: the original procedure is designed to surface genes whose patient–cell-line discrepancy is statistically robust under non-Gaussian distributions, and a simple correlation cannot recover this signal.

## D Biomni real-world usage analysis

We analyzed 32,014 user queries collected from the open-source Biomni platform [Huang et al., 2025]. Each query was processed by an LLM-based classifier and tagged along three multi-label axes: *query type* (high-level user intent), *biology domain*, and *task category*. Data analysis is the dominant query type (63.3%), well ahead of literature research (55.0%), experimental design (26.2%), quick factual queries (25.1%), and manuscript writing (15.4%); percentages exceed 100% because 77.0% of queries carry more than one intent (Figure 5). This dominance motivates the choice of data analysis as the first release of BiomniBench. Top biology domains in the queries include cancer biology, immunology, and neuroscience; the most common analytical tasks span gene expression analysis, differential expression, statistical analysis, and pathway analysis.

**Figure 5:**
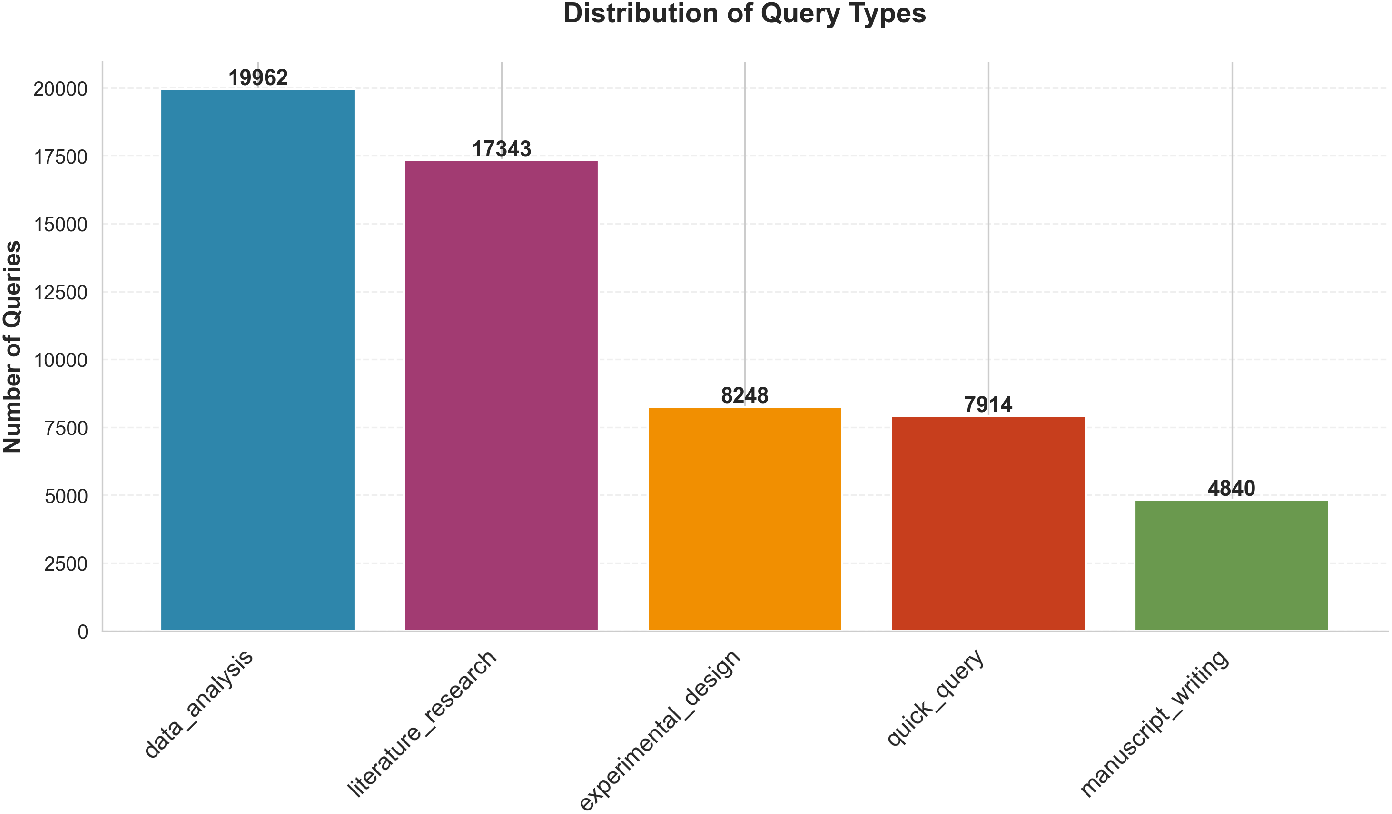
Distribution of query types across 32,014 Biomni user queries. Multi-label classification: percentages sum to greater than 100% because 77.0% of queries carry more than one type. Data analysis is the dominant intent at 63.3%.

## E Source publications

